# Proteomic and Evolutionary Analyses of Sperm Activation Identify Uncharacterized Genes in *Caenorhabditis* Nematodes

**DOI:** 10.1101/290221

**Authors:** Katja R. Kasimatis, Megan J. Moerdyk-Schauwecker, Nadine Timmermeyer, Patrick C. Phillips

## Abstract

**Background:** Nematode sperm have unique and highly diverged morphology and molecular biology. In particular, nematode sperm contain subcellular vesicles known as membranous organelles that are necessary for male fertility, yet play a still unknown role in overall sperm function. Here we take a novel proteomic approach to characterize the functional protein complement of membranous organelles in two *Caenorhabditis* species: *C. elegans* and *C. remanei*.

**Results:** We identify distinct protein compositions between membranous organelles and the activated sperm body. Two particularly interesting and undescribed gene families—the Nematode-Specific Peptide family, group D and the here designated Nematode-Specific Peptide family, group F—localize to the membranous organelle. Both multigene families are nematode-specific and exhibit patterns of conserved evolution specific to the *Caenorhabditis* clade. These data suggest gene family dynamics may be a more prevalent mode of evolution than sequence divergence within sperm. Using a CRISPR-based knock-out of the NSPF gene family, we find no evidence of a male fertility effect of these genes, despite their high protein abundance within the membranous organelles.

**Conclusions:** Our study identifies key components of this unique subcellular sperm component and establishes a path toward revealing their underlying role in reproduction.

## Background

Despite coming in a wide variety of morphologies, sperm exhibit three key cellular traits that are widely conserved across metazoans [reviewed in 1,2]. First, it appears all sperm undergo a histone-to-protamine chromatin condensation [3]. Second, the vast majority of sperm swim using a flagellum coupled to an actin/myosin cytoskeleton [4]. Third, most sperm contain an acrosome or acrosome-like membrane domain that aids in sperm-egg recognition and fusion [5]. In contrast to other animals, the phylum Nematoda has a distinctly different sperm morphology and molecular biology [6]. Namely, nematodes have large, amoeboid-like sperm cells that use non-actin mediated locomotion [7]. While other species with aflagellate sperm rely on passive diffusion for locomotion [1,4], nematodes use Major Sperm Protein (MSP)-mediated motility to crawl [7,8]. Nematode sperm also lack an acrosome [6], and membrane remodeling during spermiogenesis (sperm activation) is instead largely driven by membranous organelles [9]. Both the use of MSP-mediated motility and the presence of membranous organelles are critical components of nematode sperm biology that are unique to and conserved across this ancient phylum.

Perhaps not surprisingly, these two unique components of nematode sperm interact with one another throughout spermatogenesis. Membranous organelles are membrane bound vesicles derived from the Golgi that are found throughout the dividing cell [9]. Membranous organelles migrate to the periphery of unactivated spermatids where they associate with MSP to form fibrous body membranous organelles (Fig. 1A). During spermiogenesis MSP dissociates from the membranous organelle to form branching filaments, which structure the pseudopod of motile sperm [10,11]. Meanwhile, the membranous organelles remain associated with the cell body, fusing with the cell membrane to create cup-like structures reminiscent of secretory vesicles [8,9]. Unlike an acrosome reaction, however, the membranous organelles fuse prior to any contact with an oocyte. The role of membranous organelles and the function of these fusion events remains unknown, largely because of the challenge of studying subcellular components in single gametes. Nevertheless, mutant screens targeting faulty spermatogenesis have shown that incorrect membranous organelle fusion results in sterility {Achanzar:1997vc} and therefore that these organelles must play an important functional role within sperm. One hypothesis for membranous organelle function is that the increased membrane surface area and incorporation of additional proteins is important for membrane microdomain remodeling and fluidity [12,13]. Alternatively, since membranous organelles release their contents into the extracellular space, they may function as a source of seminal fluid proteins and therefore be involved in post-insemination reproductive tract dynamics. However, without additional information on the composition of membranous organelles, determining their functional role is a challenge.

**Figure 1.**
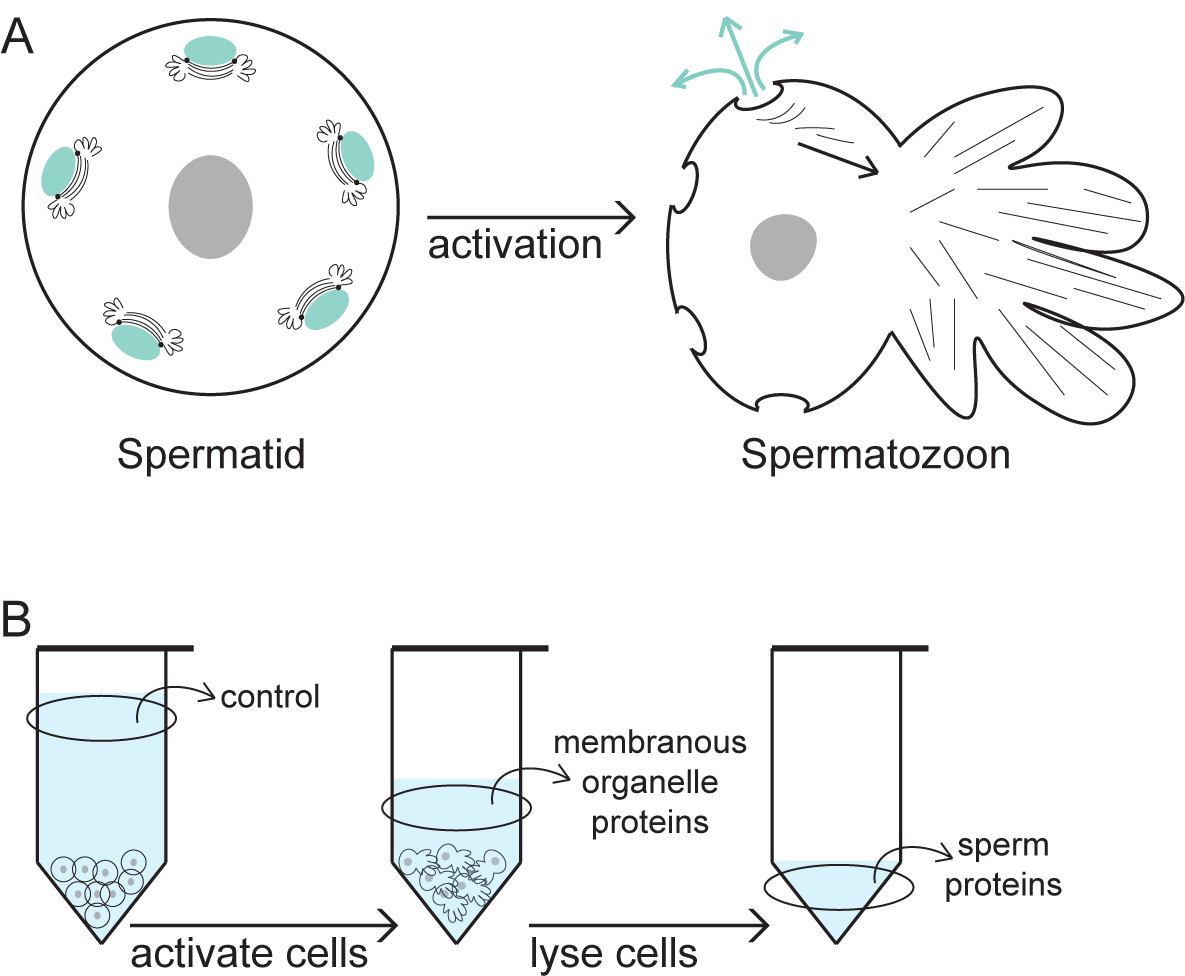
Spermiogenesis in nematodes. A) In un-activated spermatids, membranous organelles (shown in teal) form an association with Major Sperm Protein. Upon sperm activation, Major Sperm Protein forms the pseudopod of the cell and is used to crawl, while the membranous organelles fuse with the cell membrane and release their contents into the extracellular space. B) Diagram of the *in vitro* sperm activation method used. Un-activated spermatids were collected and concentrated. The supernatant before sperm activation represents a control for cell lysis. Spermatids were activated by changing the intracellular pH. The supernatant after activation represents the proteins released during membranous organelle fusion. The activated sperm cells were lysed and the membranes pelleted. The supernatant after cell lysis represents the proteins associated with the activated sperm body.

Here we take a novel approach that co-opts sperm activation events to proteomically characterize membranous organelles within two *Caenorhabditis* species. We identify two particularly interesting gene families—the Nematode-Specific Peptide family, group D and Nematode-Specific Peptide family, group F—that are previously undescribed and use evolutionary analysis and genomic knockouts to more directly probe their function.

## Results

### Proteomic characterization of spermiogenesis in *C. elegans*

Un-activated spermatids were collected from males using a novel microfluidic dissection technique coupled with a male crushing approach [modified from 14,15]. The male dissection technique utilizes a custom microfluidic device with a fine glass needle to slice through the cuticle and testis of males to release stored spermatids (Fig. 2). Alternatively, the male crushing technique squeezes the testis out of males to release spermatids. Three proteomes were characterized: un-activated spermatid, membranous organelle, and activated sperm (Fig. 1B). The un-activated spermatid proteome was dominated by the MSP, confirming that pure sperm cell samples were being collected (Additional File 1). The most abundant proteins, however, were from the Nematode-Specific Peptide family, group D (NSPD), which comprised approximately 50% of the total protein abundance. Since mass spectrometry identified a single peptide motif for these proteins, NSPD abundance was described at the gene family level. The NSPD family is uncharacterized, but has been previously shown to exhibit a pattern of male-enriched expression [16]. Actin proteins were also identified at < 1% abundance, which is comparable to previous biochemical estimates [7]. While relatively few total protein calls were made, fully one third of the un-activated spermatid proteome is previously uncharacterized in biological function.

**Figure 2.**
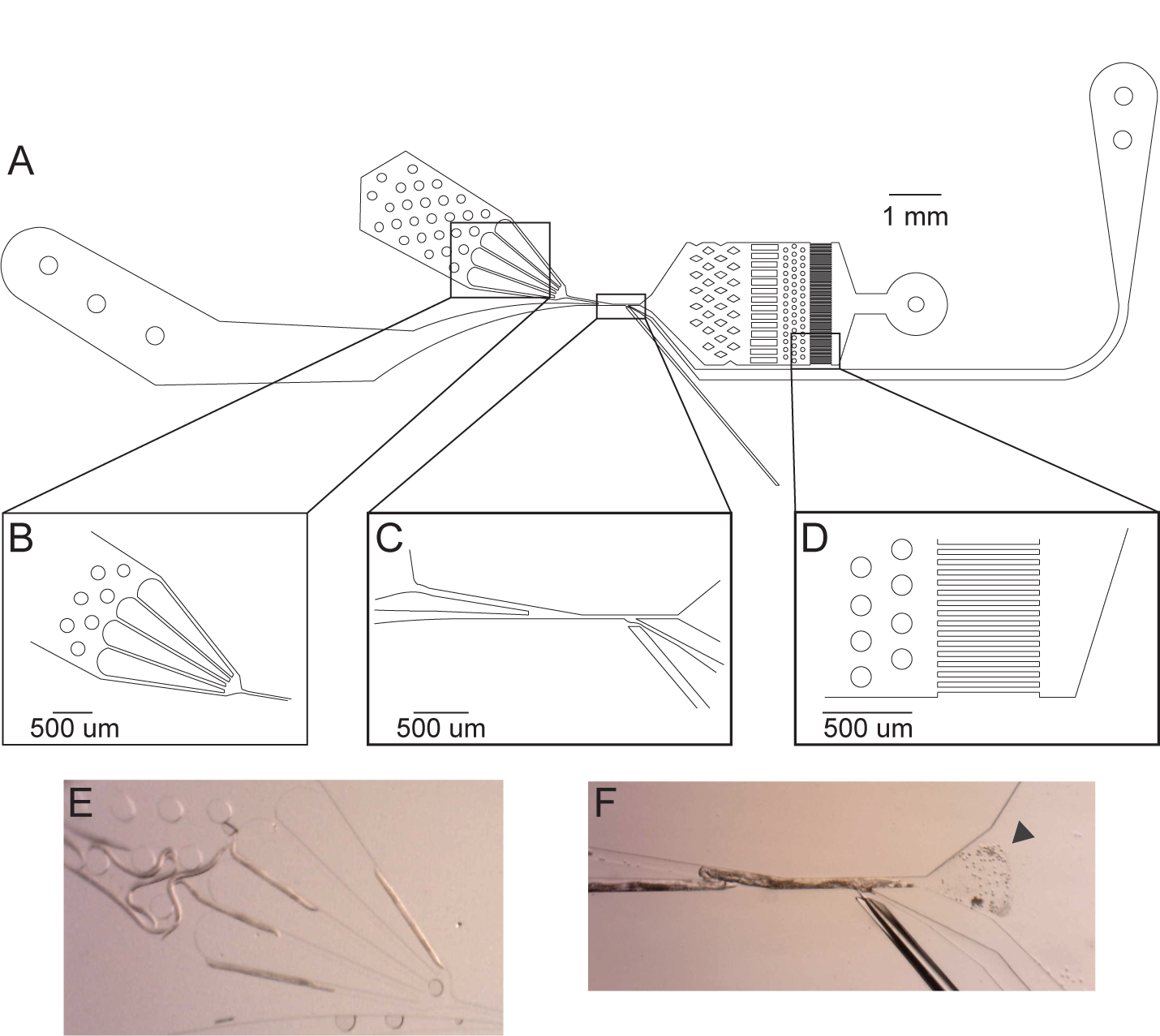
Schematic of The Shredder. A) The Shredder is a microfluidic dissection device with a single worm loading arena, a needle insertion, a sperm filtration and collection arena, and two flush channels. B) The male loading arena. The bifurcating design sequentially loads males into the dissection channel. C) The male dissection channel. Males are pushed into the channel from the loading arena and sperm cells are flushed out the right. The needle channel is separated from the male dissection channel by a thin filament of PDMS, which creates a water-tight seal around the needle. D) The sperm filter (10 um) prevents collection of non-sperm components. E) Males in the loading arena for sequential loading into the dissection channel. F) Dissected male and released spermatids (indicated by the triangle) for collection.

To isolate membranous organelle proteins from those associated with the sperm body, we took advantage of natural membranous organelle-membrane fusion during sperm activation. Spermatids were activated *in vitro* by changing the intracellular pH [9] and the proteomes of the membranous organelle secretions and activated sperm fractions were collected via centrifugation (Fig. 1B). Again, the MSP was in high abundance, though now identified in both the membranous organelle and activated sperm proteomes (Additional File 2). Interestingly, our data reveal three previously unannotated genes (Y59E9AR.7, Y59H11AM.1, and ZK1248.4) as MSPs based on high nucleotide sequence identity and presence of the MSP domain [17]. Overall, 62% of the proteins identified in either the membranous organelle or activated sperm proteome were also identified in the spermatid proteome, and all the proteins identified were previously found in the un-activated spermatid proteome collected by Ma *et al.* [18].

The proteins released from the membranous organelle during activation were distinct from those remaining in the activated sperm (Fig. 3). Seventeen proteins were unique to the membranous organelle proteome, including the NSPD family, which comprised 10% of the total membranous organelle protein abundance. The actin gene family was also unique to the membranous organelle, as were several other housekeeping-related gene families. Within the activated sperm proteome, we identified 14 unique proteins, the majority of which were involved in energy production. There were 29 proteins expressed in both proteomes. To determine if any of these shared proteins were enriched in one of the proteomes, we fit a linear regression (F_1,27_ = 4.21, p = 0.05) and identified those proteins with an abundance more than two standard deviations from the regression line (Fig. 4). The genes F34D6.7, F34D6.8, and F34D6.9 were identified as significantly enriched in the membranous organelle proteome and were in fact the most abundant membranous organelle protein after MSP. Again, these protein calls were described using a single abundance measure due to identical mass spectrometry peptide sequence identification. The F34D6 parent sequence describes genes in the 2,680 kb region of Chromosome II in the *C. elegans* genome. The three F34D6 genes identified in the proteomic data are distinct from other genes in this region, uncharacterized, and display male-specific expression [16], consistent with our observations.

**Figure 3.**
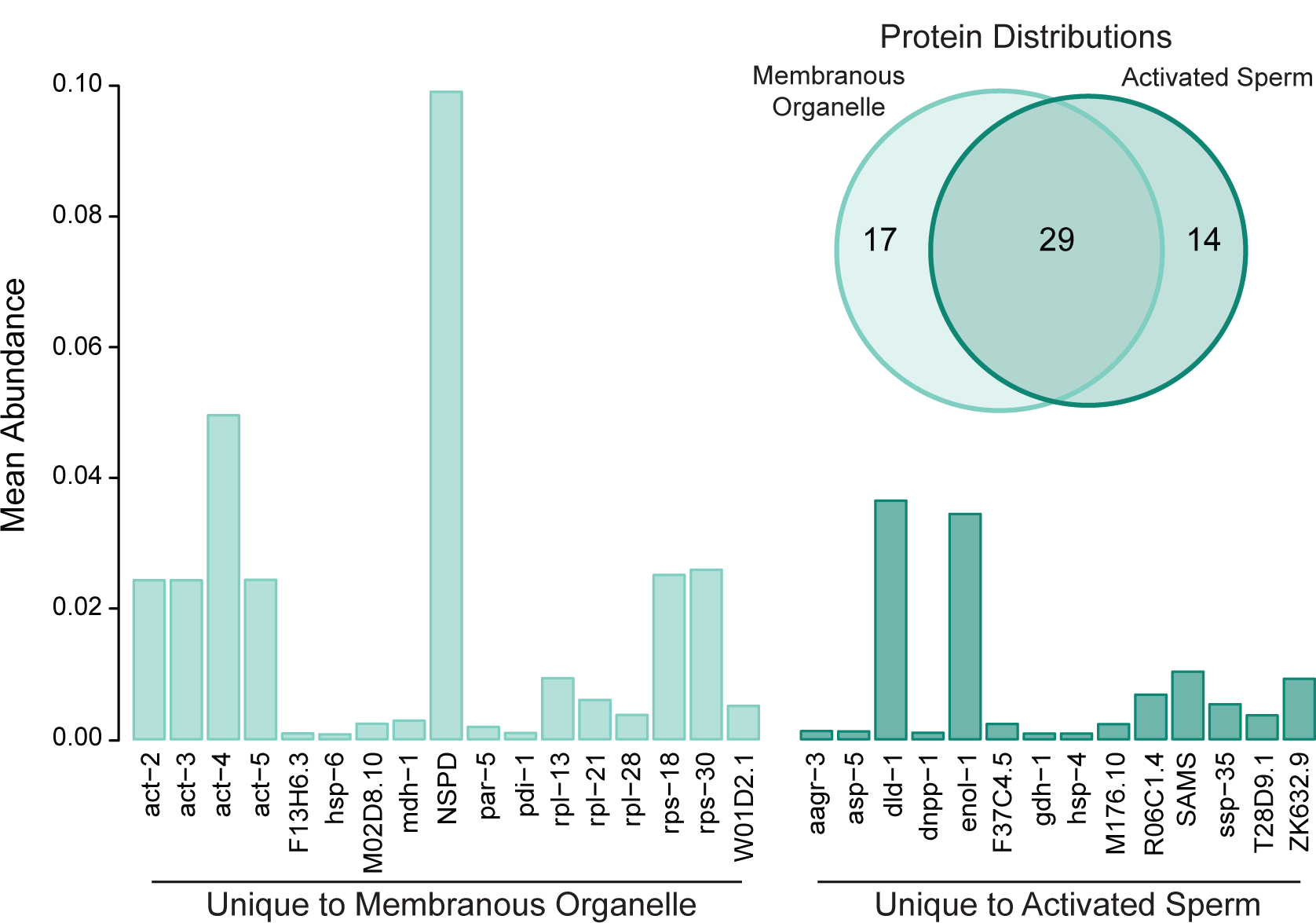
Proteomic characterization of the membranous organelle and activated sperm proteomes in *C. elegans*. The two proteomes were distinct, with 17 proteins found only in membranous organelles and 14 proteins found only in activated sperm. The proteins unique to the membranous organelles include the Nematode-Specific Peptide family, group D (NSPD) as well as several housekeeping gene families. The proteins unique to activated sperm are predominantly involved in energy production. Protein abundance is shown as the mean normalized spectrum abundance frequency.

**Figure 4.**
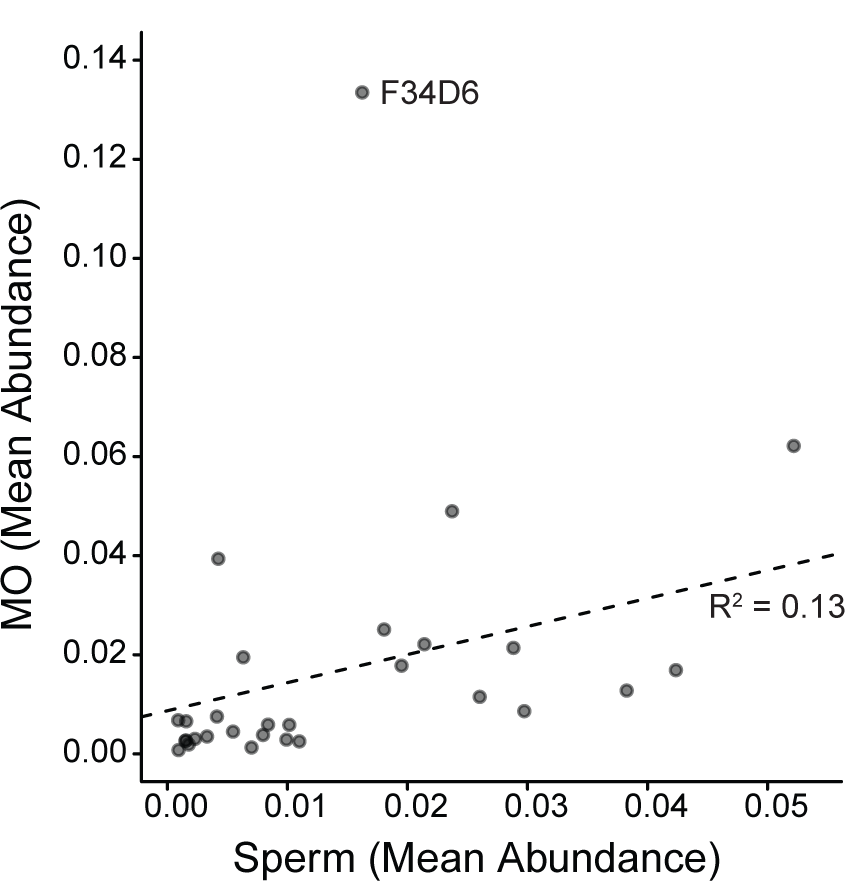
The 29 proteins shared between the activated sperm and membranous organelle in *C. elegans*. A linear regression between the two subsets (F_1, 27_ = 4.21, p = 0.05) shows the F34D6 protein to be highly enriched in the membranous organelle. Protein abundance is shown as the mean normalized spectrum abundance frequency. Here, the Major Sperm Protein was excluded from this analysis as an outlier, although the results remain qualitatively similar if it is included.

### Proteome composition is largely conserved between species

Spermatids were also collected from the obligate outcrossing nematode *C. remanei.* To compare proteome composition between divergent species, we condensed all protein calls to the gene family level. Within *C. remanei*, we identified 64 gene families in the membranous organelle proteome and 94 gene families within the activated sperm proteome, with 51 families being shared between the proteomes (Additional File 2). Of all the proteins identified, nine did not have an annotated *C. elegans* ortholog. A total of 34 gene families were identified in both *C. elegans* and *C. remanei*, capturing the majority of highly abundant genes identified. However, more proteins of low abundance were identified in *C. remanei*. Three gene families – NSPD, Actin, and Ribosomal Proteins, Large subunit – unique to the membranous organelle proteome in *C. elegans* were identified in low abundance within activated sperm in *C. remanei*, potentially because of differential success in activating *C. remanei* sperm *in vitro* (Additional File 2). Two noticeable differences between species were the presence of histone proteins and the absence of F34D6 orthologs in *C. remanei*.

### Evolutionary analysis of membranous organelle proteins

Proteomic analysis identified NSPD and F34D6 proteins as being highly abundant and localized their expression to the membranous organelle. Yet no information exists about the molecular or biological function of these genes. To better understand the nature of these gene families, we analyzed their evolutionary history across the *Elegans* supergroup within *Caenorhabditis*. We made custom annotations of these gene families in 11 species using the annotated *C. elegans* genes (ten NSPD and F34D6.7, F34D6.8, and F34D6.9) as the query dataset. Our sampling included the three lineage transitions to self-fertilizing hermaphroditism and the single lineage transition to sperm gigantism found within this clade.

Across all 12 species we identified 69 NSPD homologs (Additional File 3). The NSPD gene family ranged from three to ten gene copies, with *C. elegans* having the highest copy number and *C. kamaaina* having the fewest (Fig. 5). Coding sequence length was largely conserved between paralogs, but differed across species. Sequence length differences were particularly driven by a 24-30 base pair region in the middle of the gene containing repeating of asparagine and glycine amino acids, which tended to be the same length within a species, but differed across species. Despite these species-specific repeats, amino acid sequence identity between paralogs was high, ranging from 81.3-95.3%. No secondary structure was predicted for these genes and in fact they were biochemically categorized as being 73% intrinsically disordered due to low sequence complexity and amino acid composition biases [19,20]. The NSPD genes were broadly distributed across the genome, occurring as single copies on multiple chromosomes or scaffolds in each species (Additional File 3). This seemingly independent arrangement of individual genes throughout the genome precluded a robust syntentic analysis. Additionally, phylogenetic analysis showed NSPD genes predominantly cluster within species and thus they do not convey a strong signal of ancestral gene orthology (Additional File 4). Since orthologous genes could not be assigned, the protein coding sequences were analyzed within the four monophyletic clades represented. Even within these shorter evolutionary timescales, orthologous genes were not readily apparent, again suggesting species-specific evolution at the gene family level. To assess variation in evolutionary rate across the gene family, we estimated a single, alignment-wide ratio of non-synonymous to synonymous substitutions (ω) using reduced sequence alignments. Specifically, we removed the species-specific amino acid repeats in the middle of the gene, which were highly sensitive to alignment parameters. The ω-values varied widely from 0.07 to 0.37 with the more recently derived clades having higher values (Fig. 5), although none indicate a strong signal of positive selection.

**Figure 5.**
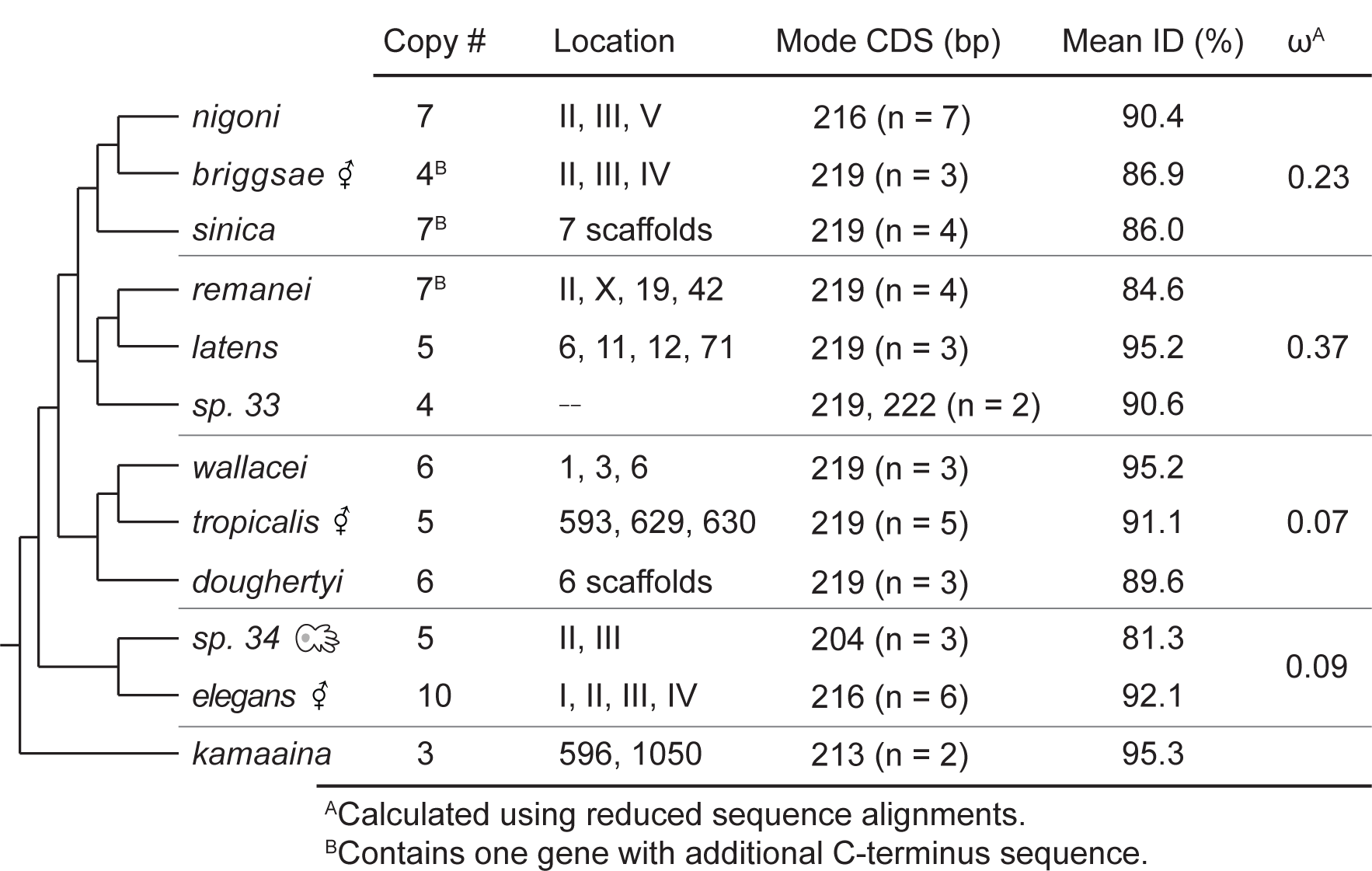
The evolution of the Nematode-Specific Peptide family, group D (NSPD) across the *Caenorhabditis Elegans* Supergroup. Listed for each species are: the number of gene copies annotated, the genomic location (Roman numerals represent chromosome level assemblies and numbers represent scaffolds), the mode coding sequence length in base pairs (n = number of gene copies of said length), the mean amino acid sequence identity between paralogs, and the alignment-wide estimate of the ratio of non-synonymous to synonymous substitutions (ω). The complete gene annotation list is provided in Additional File 3.

The F34D6.7, F34D6.8, and F34D6.9 genes in *C. elegans*, as uniquely identified within the membranous organelle fraction of the sperm proteome, are organized as an array and have a nucleotide sequence similarity of 93.9%. Given their genomic organization, sequence similarity, and co-localization of expression, these genes appear to be a small gene family that originated via tandem duplication. Additionally, an amino acid blast search of these F34D6 sequences in NCBI reveals that they are nematode-specific. Thus, they comprise a newly identified Nematode-Specific Peptide family, which we designate as NSP group F (NSPF). Like the NSPD family, these genes do not have a predicted secondary structure and are 40% intrinsically disordered. They are, however, biochemically predicted to be signaling peptides (mean signal peptide score = 0.9) with a predicted cleavage site between amino acid residues 19 and 20.

We identified and annotated 22 NSPF orthologs in ten species (Additional File 3). No genes were located within *C. sp. 34* genome (which is very well assembled). Nine species had two gene copies, while *C. doughertyi* has a single copy and, as mentioned, *C. elegans* has three annotated copies. Examination of 249 sequenced *C. elegans* natural isolates [21] suggests that *nspf-2* arose through a duplication of *nspf-1*. In particular, the start codon for *nspf-1* aligns to the same position across isolates, but does not for *nspf-2*. This duplication appears fixed within the *C. elegans* lineage––though one strain (CB4856) has a premature stop codon–– and sequence identity is high between duplicates. Additionally, the *C. elegans* NSPF gene family has translocated to Chromosome II while the other species show conserved synteny to Chromosome IV (Fig. 6). Using syntenic relationships coupled with gene orientation and phylogenetic clustering, we were able to assign gene orthology within the family (Additional File 5). Within these orthologous groups, species relationships were largely recapitulated with ω-values of 0.53 and 0.26 for the *nspf-1* and *nspf-3* orthologs, respectively. However, when the *C. elegans* lineage was excluded, the ω-values sharply decreased to 0.15 for the *nspf-1* and 0.17 for the *nspf-3* orthologs, indicating a pattern of sequence constraint (Fig. 6). We explicitly tested if the *C. elegans* lineage was evolving at a different rate than the other lineages. Indeed, the *nspf-1* (ω = 1.1, C.I. of ω = 0.78 – 1.5, -2Δ*ln* = 5.11) and to a lesser extent the *nspf-3* (ω = 0.57, C.I. of ω = 0.34 – 0.87, -2Δ*ln* = 2.34) *C. elegans* lineages showed some evidence of positive selection, although the differences in the likelihoods of the two models were not statistical significant.

**Figure 6.**
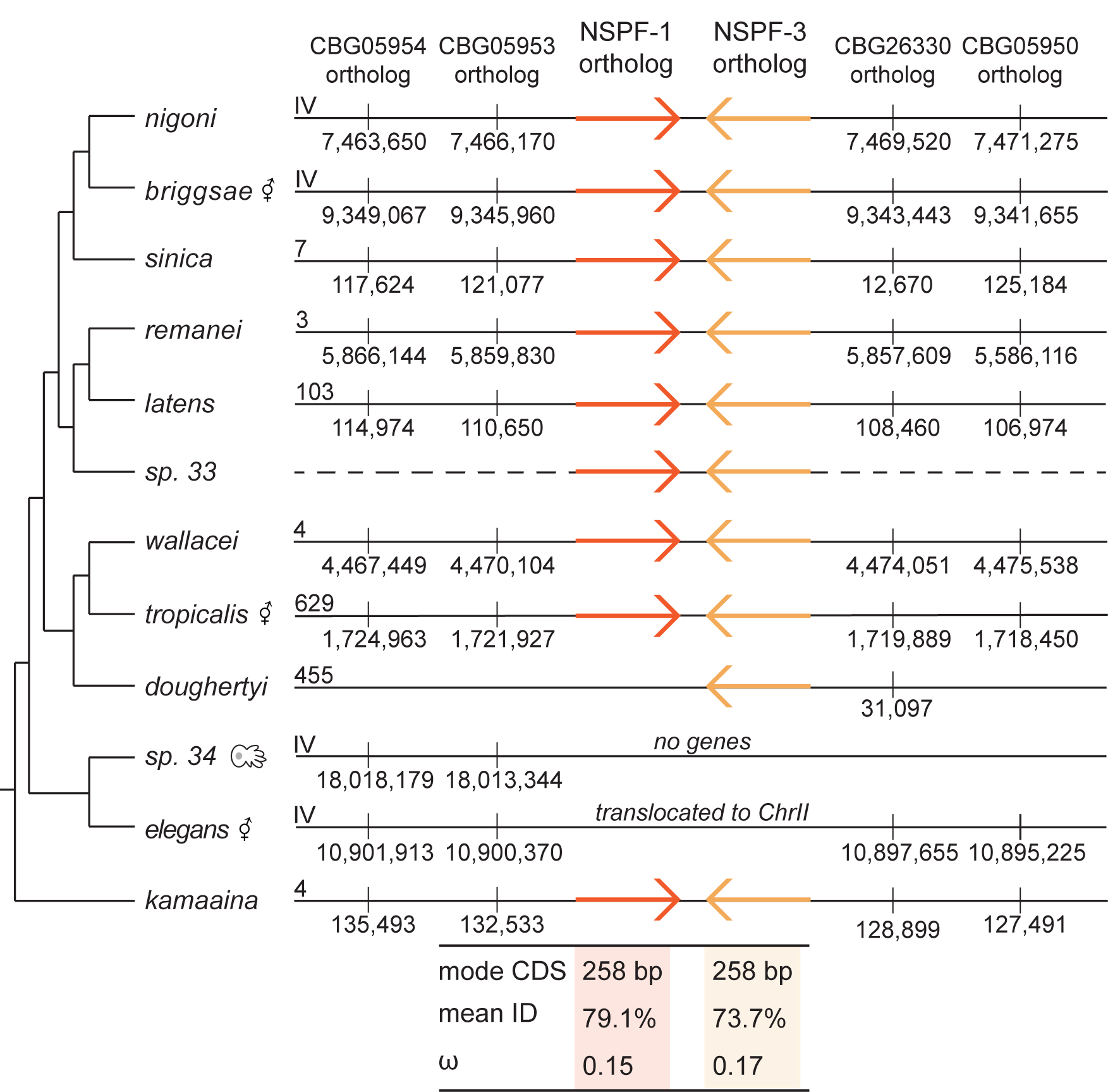
The evolution of the NSPF gene family across the *Elegans* Supergroup. The orthologous *nspf-1* and *nspf-3* genes are shown in orange on the chromosome or scaffold to which they locate. The Chromosome IV gene anchors used to determine synteny are shown. For each orthologous group the mode coding sequence length (in base pairs), the mean amino acid sequence identity, and the alignment-wide estimate of the ratio of non-synonymous to synonymous substitutions (ω) are shown. The *C. elegans* orthologs are excluded from the mean identity and ω estimates as they show distinctly different patterns of evolution. The complete gene annotation list is provided in Additional File 3.

### Functional analysis of the NSPF gene family

Given the high abundance of the NSPF protein, the conserved nature of these genes, and their potential as signaling peptides, we hypothesized these genes could be important for male fertility either during spermatogenesis or in sperm competition. Using CRISPR, we knocked out the three NSPF genes in the *C. elegans* standard laboratory strain (N2) to directly test the function of this gene family. We quantified male reproductive success, by allowing single males to mate with an excess of females over a 24 hour period. Very little difference in progeny production was observed between knockout and wildtype males (t = -0.81, df = 26.0, p = 0.42; Fig. 7A). Given the size of our experiment and the large sampling variance in individual fecundity, we would have been able to detect a difference between backgrounds of 24% with 80% power, so we possibly missed some effects if they were particularly subtle. We also measured the role of these genes in male competitive success, finding again that knocking out these genes had no effect on male fertility (Fig. 7B). In fact, knockout males were no worse competitors than wildtype males (z = -0.12, p = 0.90) and produced roughly 50% of the progeny measured (proportions test: χ^2^ = 1.27, df = 1, p = 0.26, C.I. of progeny produced = 27.4 – 55.9%). Overall, then, despite is prevalence within the sperm membranous organelle, the NSPF gene family does not appear to play an important role in male fertilization success.

**Figure 7.**
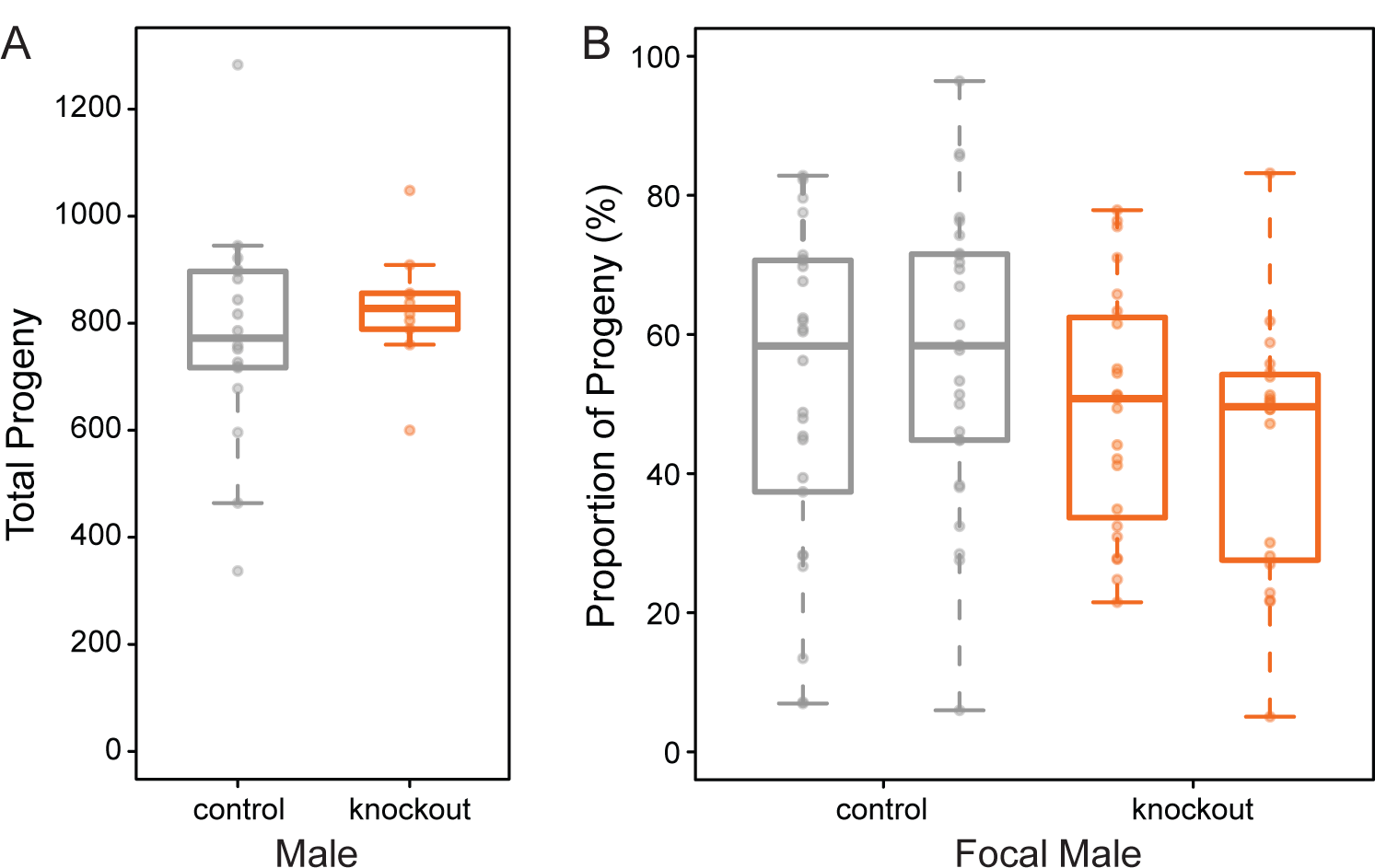
Functional assays of the NSPF gene family in *C. elegans* male fertility. A) In a non-competitive sperm setting, knockout males (orange) do not produce significantly fewer progeny than control males (gray) when given an excess of females with which to mate (t = -0.81, df = 26.0, p = 0.42). B) In a competitive sperm setting, knockout males (orange) do not produce significantly fewer progeny than control wildtype males (z = -0.12, p = 0.90) nor do they produce a significant deviation from 50% of the total progeny production (proportions test: χ^2^ = 1.27, df = 1, p = 0.26, C.I. of progeny produced = 27.4 – 55.9%). All fecundity data are provided in Additional File 7.

## Discussion

We used a proteomic approach coupled with molecular evolution analyses and direct functional assays to characterize the composition and role of membranous organelles in nematode sperm. Our approach capitalized upon the natural sperm activation process to accurately isolate membranous organelle proteins for the first time. This proteome set captures the most abundant proteins found in sperm and shows that the composition of the membranous organelle proteome is distinct from that of the activated sperm body. Unsurprisingly, the most abundant sperm protein was the major sperm protein (MSP). Interestingly, MSPs were also the most abundant proteins in the membranous organelle. Since MSP proteins are important not only for motility, but also for oocyte signaling [22], identifying them as an abundant membranous organelle component implicates membranous organelle fusion as a method by which free-floating MSP is added to the seminal fluid. There are 31 annotated MSP gene copies in *C. elegans*, with potentially more uncharacterized copies as seen here, and as of yet we do not know if some of them might be subfunctionally located within different parts of the sperm [17]. We also found that sperm proteome composition was largely conserved between *C. elegans* and *C. remanei*, particularly within the activated sperm itself. This is the first investigation of the proteome of a gonochoristic nematode. Although similarity is the rule, we did identify several *C. remanei* proteins lacking *C. elegans* orthologs, which are potentially unique and warrant future molecular characterization.

Two gene families identified in the membranous organelles are particularly notable. First, the NSPD gene family was unique to the membranous organelle. This previously uncharacterized gene family shows high sequence similarity between paralogs and low levels of divergence between species. The high degree of similarity between paralogs is particularly interesting as these genes are not organized as a single cluster and therefore sequence similarity is likely not maintained through non-homologous DNA repair (i.e., gene conversion) [23]. Additionally, NSPDs lack secondary structure and are in fact predicted to be intrinsically disordered. This lack of divergence coupled with little biochemical constraint is unusual and suggests NSPD function requires a specific amino acid sequence along its entire length. However, not all regions of the gene appear to be under the same constraint, as evidenced by the short species-specific repeating motif, although the functional relevance of this motif remains unknown. The pattern of seemingly independent gene copy number expansion and genomic organization despite sequence constraint observed here is strikingly similar to the evolutionary pattern we previously observed in the MSP gene family [17], and suggests lineage-specific gene family evolution rather than preservation of an ancestral gene family structure.

Similarly, the newly defined NSPF family has enriched expression in the membranous organelle, as well as sequence conservation across the clade. While the degree of gene family evolution was far more limited, the duplication of *nspf-2* in *C. elegans* isolates combined with apparent gene losses in *C. sp. 34* and *C. doughertyi* suggest that this family is not particularly static. The *C. elegans* lineage, in particular, appears to be evolving differently from the rest of the genus, including changes in copy number and genomic organization. Despite their predicted signaling function, we found no compelling evidence that these genes are involved in male reproductive success, though a subtle fertility difference could have been swamped out by the high individual variance in fecundity. These null results suggest that this family could be redundant as is supported by apparent species-specific gene losses, although if true we might expect to see greater sequence divergence across the genus due to genetic drift. Alternatively, this family may play a role in female post-mating physiological response or male re-mating behavior and not on male fertility *per se*.

While these data represent a foundation for membranous organelle molecular biology, no clear functional role for this subcellular component stands out. Nevertheless, two non-exclusive hypotheses suggest themselves. First, membranous organelles may serve as a contributor to the overall composition of the seminal fluid (although perhaps a minor contributor). The presence of MSP within the organelles supports this hypothesis. Future studies that track where membranous organelle proteins are found after activation—in the male vas deferens, at the female vulva opening, or in the spermatheca—will be valuable in verifying this hypothesis.

Alternatively, the membranous organelle could be more important during spermatid stasis and establishing membrane fluidity upon activation [13,24]. Here, membrane fusion is the more critical functional component, and the release of membranous organelle contents would then represent an incidental “trash dump” as sperm cells move on to the next phase of their life cycle. The presence of actin exclusively in the membranous organelle supports this hypothesis, as activated sperm function is known to be actin-independent. Both hypotheses warrant continued investigation to further understand the functional role of this unique sperm component.

## Conclusions

Overall, our findings of sequence conservation over such long evolutionary time periods are contrary to observations within many other organisms, where elevated signals of positive selection are detected in seminal fluid proteins [25-27]. From an evolutionary perspective, then, patterns of evolution in secreted membranous organelle proteins do not match expectations for typical seminal fluid proteins. However, this pattern of sequence conservation coupled with lineage-specific gene family evolution observed here has also been previously identified for the MSP gene family [17]. There thus appears to be a “nematode sperm protein evolution syndrome” in which structural rearrangements and copy number variants are a more prevalent mechanism of genetic evolution than sequence divergence *per se*. Such a pattern could potentially be due to the conserved and unique sperm biology in nematodes, especially the biochemistry of locomotion. These results further support the need for taking a holistic approach when understanding the evolutionary history of genes.

## Methods

### Sperm collection

#### Worm culture and strains

Sperm were collected from *Caenorhabditis elegans* (standard laboratory strain N2 and strain JK574: *fog-2*(q71) V on the N2 background) and *C. remanei* (strain EM464). The *fog-2* mutation blocks *C. elegans* hermaphrodite self-sperm production, resulting in a functionally male-female population, thereby increasing the ease with which males could be collected. All strains were raised on NGM-agar plates seeded with OP50 *Escherichia coli* bacteria and raised at 20ºC [28]. Synchronized cultures of larval stage 1 animals were produced through hypochlorite treatment [29]. Males sourced for microfluidic dissection were isolated from females starting as young adults (44 hours post-larval stage 1) for 24 hours to build up their stored spermatid supply. Males sourced for testis crushing were maintained on mixed sex plates at population densities of approximately 1,000 animals until the second day of adulthood (62 hours post-larval stage 1).

#### Microfluidic-based sperm collection

The Shredder (final design: v5.0; Additional File 6) was designed using CAD software (Vectorworks 2013 SP5, Nemetschek Vectorworks, Inc) to function as a precise method of dissecting the male testis. The design has a single worm inlet that sequentially pushes males past a glass dissection needle, which slices through the cuticle, punctures the testis, and releases stored spermatids (Fig. 2). Two additional liquid channels flush males out of the dissection channel and flush sperm through a filtration system into the sperm outlet. Single layer devices were fabricated from polydimethylsiloxane (PDMS) using soft lithography [30] and bonded to a glass microscopy slide following exposure to air plasma. Dissection needles were made using a laser micropipette puller (Sutter Instrument P-2000) and inserted into each device following bonding.

A single Shredder could be used once to dissect up to 20 males. Each device was first flushed with 20 mM ammonium bicarbonate (pH 7.8), after which 20 virgin males were loaded into the worm inlet. The collected spermatids were concentrated by centrifugation (500 rcf for 15 minutes) and then lysed in liquid nitrogen. The cell membranes were pelleted, leaving the spermatid proteins in the supernatant for collection. A total of four pooled *C. elegans* replicates (259 males) and five pooled *C. remanei* replicates (265 males) formed the un-activated spermatid proteome for each species.

#### Testis-crushing sperm collection

To increase throughput, we also used a male crushing technique to collect spermatids [modified from 14,15]. Males were raised in mixed sex populations and size separated from females on the second day of adulthood. This developmental time point was optimal for maximizing the difference in diameter between the sexes and minimizing progeny. The sexes were separated using Nitex nylon filters (35 um grid for *C. elegans* and 30 um grid for *C. remanei*) with an average male purity of 91%. The filtration set-up was kept within a sterilized box to reduce external contamination.

Males were pelleted and plated between two 6” x 6”, silane-coated (tridecafluoro-1,1,2,2-tetrahydrooctle-1-trichlorosilane) plexiglass squares. The plexiglass was then placed between two 6” x 6” x 1” wooden blocks. A heavy-duty bench vise was used to apply pressure to males, releasing the testis and spermatids. Spermatids were washed off the plexiglass using 20 mM ammonium bicarbonate (pH 5.6) onto a 10 um grid Nitex nylon filter. This filter size was large enough to let spermatids freely pass, but not adult carcasses or eggs. Spermatids were concentrated by centrifugation and the supernatant collected (Fig. 1B). Supernatant collected before sperm activation was used to control for proteins released by cell lysis. No protein was measured in the pre-sperm activation supernatant. Spermatids were activated *in vitro* by adding 100 uL of 70mM triethanolamine (TEA) to the pelleted volume [9] and were left to activate on a chilled block for 15 minutes. The supernatant was collected to provide the membranous organelle proteome (Fig. 1B). The remaining activated cells were lysed as before and the proteins were collected as the activated sperm proteome. Six pooled replicates for *C. elegans* (maximum 19,075 males) and four pooled replicates for *C. remanei* (maximum 13,400 males) formed the membranous organelle and activated sperm proteomes for each species.

### Proteomic characterization of sperm

#### Tandem mass spectrometry

The proteomes were prepared and characterized by the Genome Science Mass Spectrometry Center at the University of Washington. Samples were denatured and digested according to standard protocols [31] and then analyzed on a Thermo Velos-Pro mass spectrometer coupled with a Thermo Easy nano-LC. Analytical replicates were run for each sample. MS/MS data were analyzed using the Comet database search algorithm [32] with either the *C. elegans* (PRJNA13758) or *C. remanei* (PRJNA53967) reference protein database. Peptide q-values and posterior error probabilities were calculated using Percolator [33]. Peptides were assembled into protein identification using ID picker [34] with a 1% false discovery rate cutoff.

#### Proteomic data analysis

Raw MS/MS information for each proteome was processed so as to include the minimum number of proteins that account for the observed peptides (i.e. parsimonious proteins) and filtered to exclude non-nematode proteins. Additionally, we combined isoform calls into a single gene and condensed four classes of genes (MSP family, NSPD family, SAMS family, F34D6 family) to the gene family level because of identical peptide coverage and high overall sequence similarity of paralogs. Overall, then, our final datasets were the most conservative representation of our data. We then calculated the relative normalized spectrum abundance frequency (measured NSAF divided by the total worm NSAF) for each protein. The two runs were combined by taking the mean relative NSAF of each protein.

Biological functions for each protein were assigned using WormBase when possible [16]. The composition of the membranous organelle and activated sperm proteomes were compared to determine which proteins were shared and which were unique to a given proteome. Within *C. elegans*, we looked for enrichment within the shared proteins by fitting a linear model of membranous organelle protein abundance against activated sperm protein abundance and looking for genes two or more standard deviations from the regression line. Since the *C. remanei* genome is not as well functionally annotated, *C. elegans* orthologous gene families were assigned to characterize biological function. Proteome composition between species was compared at the gene family level. All statistical analyses were performed using the R statistical language [35].

### Evolutionary analysis of the membranous organelle

#### Gene annotations

We used the well-annotated *C. elegans* reference genome (PRJNA13758: CEGMA: 100% complete, 0% partial; BUSCO 98% complete, n = 982) to compile our query dataset for the NSPD and NSPF (genes F34D6.7, F34D6.8, and F34D6.9) gene families. Genes were annotated in 11 species across the *Caenorhabditis Elegans* supergroup: *C. sp. 33* (from J. Wang), *C. sp. 34* (PRJDB5687), *C. briggsae* (PRJNA10731), *C. doughertyi* (PRJEB11002), *C. kamaaina* (QG2077_v1), *C. latens* (PX534_v1), *C. nigoni* (PRJNA384657), *C. remanei* (PRJNA248909), *C. sinica* (PRJNA194557), *C. tropicalis* (PRJNA53597), and *C. wallacei* (from E. Schwarz). Annotations were generated using custom amino acid blast (tblastn) searches in Geneious v10.2.3 [36]. Blast results were hand-curated for accuracy. In particular, five NSPF sequence motifs found to be conserved between *C. elegans* and *C. briggsae* were used as markers during annotation. We annotated a total of 59 NSPD genes and 19 NSPF family genes (Additional File 3) in the 11 species.

The *Caenorhabditis* Natural Diversity Resource [21] was used to probe the duplication and translocation of the NSPF family across the 249 isotypes identified from whole genome sequencing of 429 natural isolates. The NSPF gene region (II: 2,687,625 – 2,690,180) was extracted using SAMTOOLS. Coverage was calculated and those positions with less and 3x coverage were masked. A consensus sequence for each isotype was created. These sequences were aligned using ClustalW [37] in Geneious.

Synteny of the NSPF family was analyzed to determine gene orthology. The *C. elegans* NSPF family formed a cluster on Chromosome II, however, the *C. briggsae* NSPF family formed a cluster on Chromosome IV. Therefore, additional genes surrounding both the *C. elegans* and *C. briggsae* clusters were identified using the UCSC Genome Browser [36] to serve as syntenic Chromosome II and IV anchors, respectively, following the approach outlined in Kasimatis and Phillips [17]. The NSPD family was spread across more than half the chromosomes in *C. elegans* and *C. briggsae*, precluding rigorous syntenic analysis.

Secondary structure was predicted using the Phyre^2^ server [38]. Biochemical predications about protein structure and function were made using the Predictors of Natural Disordered Regions Server [19] and the SignalP Server [39].

#### Evolutionary rate tests

The gene sequences for the NSPF and NSPD families were aligned using ClustalW. Amino acid sequence identity was calculated for all pairwise gene combinations within a species as well as across the clade. Unrooted maximum likelihood phylogenies were constructed in PhyML [40] of orthologous genes for the NSPF family. Since orthology could not be assigned within the NSPD family, phylogenies were constructed based on monophyletic species trios. Alignment-wide estimates of the non-synonymous to synonymous substitution ratio (ω-ratio) were calculated using HyPhy [41] under a GTR mutation model. Selection within the NSPF family was estimated across the genus for orthologous genes. Additionally, orthologous genes were analyzed using a branch-site framework in the package BS-REL [42] within HyPhy to determine if the *C. elegans* branch in particular was evolving differently than the rest of the gene family. The NSPD family was analyzed using reduced alignments of all genes within monophyletic species triplets. Reduced alignments were constructed by removing the species-specific repeating amino acid motifs (~8 residues) in the middle of the gene. Here sequence alignment was highly dependent on the gap/extension penalty, thereby potentially confounding evolutionary inference.

### Functional verification of NSPF gene family

#### Strain generation by CRISPR/Cas9

Guide sequences were chosen using the CRISPRdirect [43], MIT CRISPR Design (http://crispr.mit.edu) and Sequence Scan for CRISPR [44] tools. For deletion of the *nspf-1*, *nspf-2*, and *nspf-3* genes, cr:tracrRNAs (Synthego) targeting the sequences CAGAGCCCATAATTCAAAGA<u>CGG</u> and AGATGAGATTCTAATCAGGT<u>AGG</u> were annealed and pre-incubated with Cas9 (PNA Bio) in accordance with the manufacturer protocol. Young adult N2 individuals were injected in the gonad with a final mix consisting of 1.7 μM of each cr:tracrRNA, 1.65 μg/μl Cas9 and 50 ng/μl of the oligonucleotide repair template (5’-GTAAGAATACAATTTTTCTTTGTGACTTACCGTCTGGTAGGGTGGCAGATCAGTGTTCAGAAGGAAGTGA-3’), along with an additional cr:tracrRNA and oligonucleotide repair template to allow for screening by *dpy-10* co-conversion [see 45]. Individuals from broods containing roller or dumpy individuals were screened for the deletion by PCR and confirmed by Sanger sequencing. Individuals with confirmed deletions were then crossed to males with the *him-5* mutation (strain CB4088: *him-5*(e1490) on the N2 background). The *him-5* mutation increases the frequency of X chromosome non-disjunction events during meiosis, resulting in roughly 30% male progeny from self-fertilizing hermaphrodites [46]. Five generation of backcrossing were done to purge potential off-target CRISPR affects. The resulting strain, PX623, (fx*Df*1 II; *him-5*(e1490) V) was used for functional analyses of the NSPF genes.

#### Fertility assays

We assayed the fertility of knockout males in both non-competitive and competitive sperm environments. To assess the overall reproductive success of knockout males, we mated a single knockout male with three wildtype, virgin females (strain JK574) for 24 hours. As a control, wildtype males (strain JK574) were mated to wildtype females following the same male to female ratio. Matings were done on small NGM-agar plates (35 mm diameter) seeded with 10 uL OP50 *E. coli*. After 24 hours, each male was removed and the females were transferred to a new plate to continue laying eggs. Females were transferred to new plates every 24 hours until progeny production ceased. The total number of progeny was counted as a measure of each male’s reproductive success (Additional File 7). To measure competitive ability, wildtype, virgin females (strain JK574) were mated with a knockout male and an RFP marked male. Again as a control, virgin females were mated to a wildtype male and an RFP marked male. Worms were mated overnight on small NGM-agar plates seeded with 10 uL OP50 *E. coli* and then the males were removed. Progeny were collected over the next 24 hours, counted, and screened for the number of RFP positive progeny. Two independent biological replicates of the competitive assay were performed (Additional File 7).

The fertility data were analyzed using R, with the significance of non-competitive reproductive success evaluated using Welch’s Two Sample t-test and an analysis of the power of the comparison computed using the package *pwr* [47]. Male sperm competitive success was analyzed using a generalized linear model framework with random effects and a Poisson distribution within the package *lme4* [48]. An equality of proportions test was performed for the competitive sperm assay to determine if wildtype and knockout males sired half of the total progeny.

## Abbreviations

MS/MS: Tandem mass spectrometry
NSAF: Normalized spectral abundance frequency
CRISPR: Clustered regularly interspaced short palindromic repeats

## Declarations

### Ethics approval and consent to participate

Not applicable

### Consent for publication

The authors all consent to this publication.

### Availability of data and materials

The datasets supporting the conclusions of this article are available in Additional Files 2 and 7. The genomes used are publically available from the following sources: WormBase (*C. elegans*, *C. briggsae*, *C. sinica*, *C. tropicalis*), the Caenorhabditis Genomes Project (*C. doughertyi*, *C. latens*, *C. kamaaina*, *C. remanei*), and NCBI (*C. sp. 34, C. nigoni*). The genome for *C. wallacei* was provided by E. Schwarz and the transcriptome for *C. sp. 33* was provided by J. Wang. Worm strains N2, JK574, CB4088, EM464, and PX623 are available from the *Caenorhabditis* Genetics Center.

### Author contributions

KRK and PCP designed the study; NT designed the original version of The Shredder, which was further refined by KRK; KRK collected samples for MS analysis; KRK analyzed the data; MMS and KRK created the strain; KRK performed fecundity analyses; KRK and PCP wrote the manuscript. All authors have read and approved the final version of this manuscript.

## Acknowledgements

Biological mass spectrometry was done through the Genome Sciences Mass Spectrometry Center at the University of Washington and we would especially like to thank Gennifer Merrihew and Michael MacCoss for assistance. Discussions with Willie Swanson regarding proteomic approaches to studying sperm function helped to set the stage for much of this work. Refinements of The Shredder designed were aided with contributions from Stephen Banse. John Johnson assisted with microfluidic sperm dissections. Matthew Rockman, Erich Schwarz, John Wang generously shared unpublished genomic data, and Anastasia Teterina assisted with the *Caenorhabditis* natural-isolate annotations. Finally, we would like to thank William Cresko, Michael Harms, and the Harms Lab Group for constructive feedback.

## Additional Files

**Additional File 1. The un-activated sperm proteome of *C. elegans*.** The majority of the proteome is comprised of the Nematode-Specific Peptide family, group D (NSPD) and the Major Sperm Protein (MSP). Low abundance proteins are outlined in the bar graph. Protein abundance is shown as the percent of the mean normalized spectrum abundance frequency.

**Additional File 2. Proteome data for *C. elegans* and *C. remanei*.** Un-activated spermatid, membranous organelle, and active sperm proteome data for both species analyzed, including WormBase gene identifiers, protein abundances, and peptide coverage.

**Additional File 3. Gene annotations for the NSPD and NSPF gene families.** Orthologous genes for the Nematode-Specific Peptide family, group D (NSPD) and Nematode-Specific Peptide family, group F (NSPF) family in 11 *Caenorhabditis* species. Annotations are listed by species, along with the gene start position and coding sequence length.

**Additional File 4. An unrooted maximum likelihood phylogeny for the Nematode-Specific Peptide family, group D (NSPD).** Genes tend to cluster within species and do not recapitulate an evolutionary history of gene orthology. Asterisks denote bootstrap values greater than 80%.

**Additional File 5. Unrooted maximum likelihood phylogenies for the Nematode-Specific Peptide family, group F (NSPF) orthologous genes.** Overall, gene trees recapitulate species relationships. Asterisks denote bootstrap values greater than 80%.

**Additional File 6. The Shredder microfluidic design.** The Shredder v5.0 is designed to dissect day 1 adult males. The blueprint is accessible using CAD software. A master height of 35um is recommended.

**Additional File 7. Functional assays of the Nematode-Specific Peptide family, group F (NSPF) gene family.** The fecundity data for total reproductive success and competitive reproductive success of NSPF knockout males.

